# PG-LLM: Benchmarking General-Purpose Language Models for Protein Variant Ranking

**DOI:** 10.64898/2026.07.27.741045

**Authors:** Rohit Arora, Leo Tianlai Chen, Melissa Du, Debora S. Marks, George M. Church

**Affiliations:** Harvard University; Capable

## Abstract

General-purpose frontier language models are being increasingly utilized for protein-design work, yet their ability to understand and evaluate variant effects remains unclear. Here, we introduce PG-LLM, a benchmark comprising 276 protein-variant prioritization tasks: 217 from ProteinGym and a temporally held-out set of 59 from recently published studies. Each task follows the same format: a language model is asked to rank a list of variant sequences given only the wild-type protein sequence and an assay description with no access to tools, multiple-sequence alignments, or protein structures. We evaluate thirteen language models and 95 published protein predictors on the same variants with the same evaluation metric. Claude Opus 5 (Max) and GPT 5.6 Sol (Max) are the best performing LLMs with Spearman correlations of *ρ* = 0.406 and 0.402 respectively. Opus 5 outperforms 49 of 95 published protein predictors, including 41 of 46 sequence-only methods, and approaches ESM2-650M at *ρ* = 0.411, but remains below the leading predictor VenusREM at *ρ* = 0.523. We observe that variant-ranking performance scales with test-time compute across GPT, Claude, and Gemini models, but gains taper before closing the gap to specialist protein predictors. To address contamination risk, we create a held-out evaluation set with 59 DMS assays from 19 studies whose scores first became public after January 2026. On this set, we observe performance and test time compute scaling trends similar to those on the 217 tasks derived from ProteinGym. PG-LLM shows that tool-free language models capture substantial protein-variant signal, outperforming many sequence-based predictors while remaining below the strongest specialized models.

## 1 Introduction

Prioritizing protein variants for functional testing and lead optimization is a central challenge in protein engineering. Current state-of-the-art methods are specialist variant-effect predictors that integrate evolutionary, structural, and protein-family information to estimate the consequences of sequence variation. General-purpose language models have shown improving performance across biological knowledge and reasoning benchmarks, suggesting that they could also support protein-variant prioritization. ^3;4^ Unlike specialist predictors, general-purpose LLMs do not depend on a dedicated protein-modeling pipeline. Instead, they can interpret an assay objective, compare variant sequences, and reason at inference time about which variants are most likely to perform well.

Deep mutational scanning (DMS) experiments measure the effect of hundreds to thousands of variants in a protein, producing maps of sequence–function relationships across phenotypes including activity, binding, expression, organismal fitness, and stability. ^5^ ProteinGym standardizes these measurements and provides a common evaluation framework for variant-effect prediction. ^6^ ProteinGym spans family-specific and alignment-based methods, ^7–11^ sequence-only pretrained models, ^1;12–15^ and systems that incorporate structural or retrieval-based information. ^2;16–19^

The ProteinGym-Hard evaluation reported in a recent Anthropic system card provided an early test of this capability on a subset of ProteinGym. ^20^ However, it only evaluated Claude-family models and did not compare them with other language models or the broader landscape of published protein predictors. This evaluation also does not include a deeper analysis of performance scaling with test-time compute, candidate-set size, and potential contamination.

Here, we introduce PG-LLM, a benchmark comparing thirteen general-purpose language models with 95 published protein predictors across all 217 ProteinGym substitution assays. Each task follows the same format: a language model receives a wild-type protein sequence, an assay description, and 50 mutant sequences and must rank them without access to multiple-sequence alignments or protein structures. We evaluate each language model at its strongest reasoning setting and analyze the effects of reasoning effort, variant-list length, output completeness, evolutionary depth, and explicit source-study recognition on score.

We find that while frontier language models recover substantial protein-variant signal in low-N settings, they do not yet match the strongest performing specialist systems. Claude Opus 5 achieves Spearman *ρ* = 0.406, surpassing the median sequence-only predictor and 49 of 95 published comparators while trailing leading protein-specific methods. Increasing test time compute for this task improves performance in every comparison. These results establish protein-variant ranking as an emerging capability of general-purpose language models and quantify the remaining headroom. We release the benchmark, evaluation sets, and available reasoning summaries at https://www.proteingymllm.com/.

### A GPT-5.6 Sol example: NusA stability

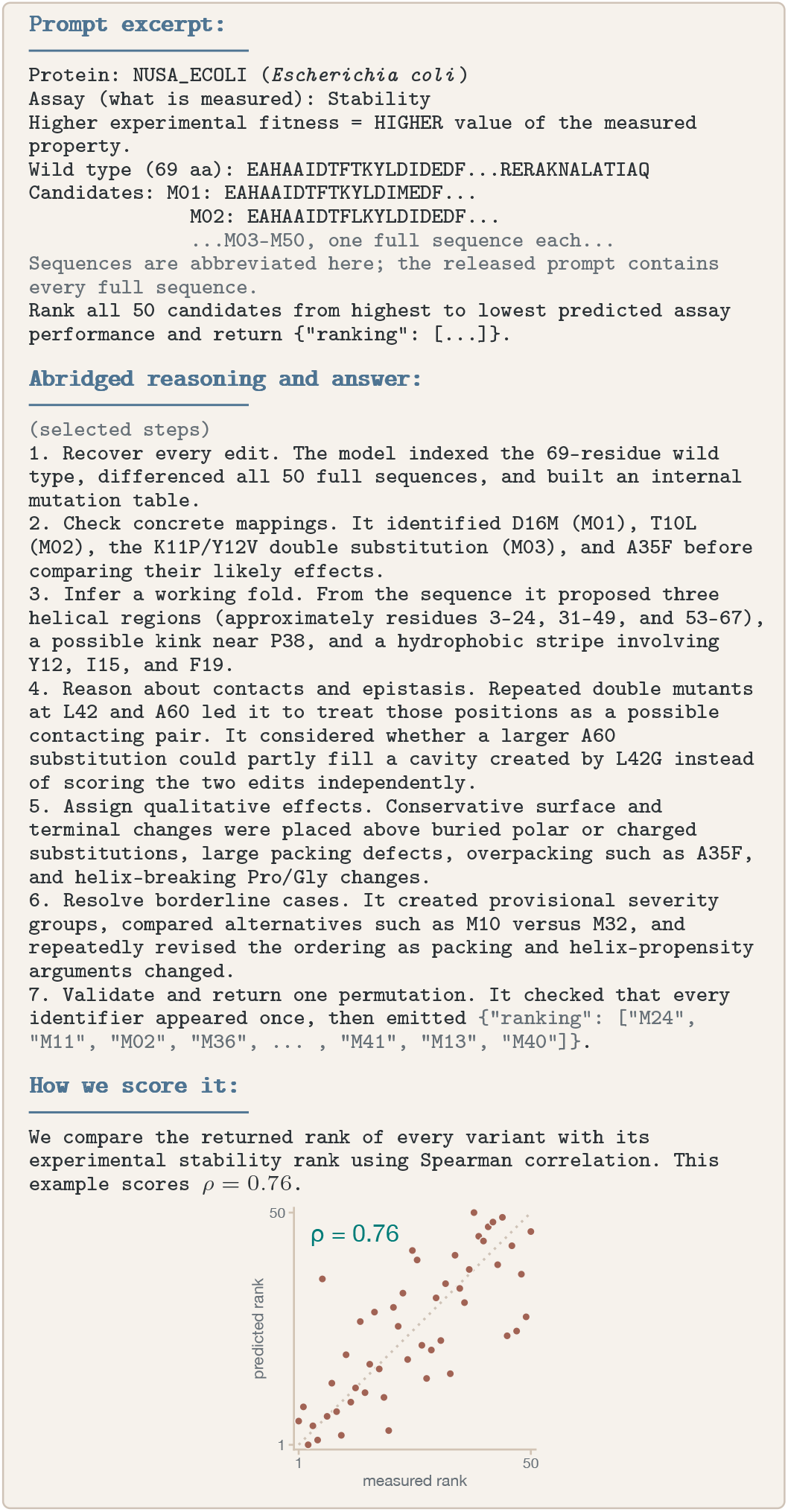

## 2 The PG-LLM benchmark

PG-LLM evaluates a single task: given an assay objective and a set of variant amino acid sequences, rank variants by their expected experimental performance. The full prompt includes the protein and organism names, a short description of the measured phenotype, the direction corresponding to better performance, the full wild-type sequence, and a shuffled set of full-length mutant sequences. The model must return one JSON ranking variants from highest to lowest predicted assay performance. The prompt does not include the source publication, multiple-sequence alignments, or structure.

### Variant-set construction

We use all 217 assays across 186 proteins in the substitution arm of ProteinGym v1.3: 148 assays contain single substitutions and 69 include multi-mutant variants. ^6^

A *variant set* is the list of variants shown in one prompt. To derive these sets, we sort variants within each assay by measured fitness, divide them into ten equally populated bins, and approximately sample the same number of variants from each bin. The resulting variant-set spans the assay’s measured fitness range. We then shuffle the variant order randomly, assign each variant a unique identifier, and display the resulting list in the prompt. If an assay contains fewer than the requested *N* variants, we use all available variants.

For each assay, we sample three independent variant sets, which we call *draws*. Draws 1–3 are generated with fixed seeds 1–3. Every LLM and published specialized predictor is evaluated on the same variants for each draw.

We also test the effect of varying variant-set lengths. In particular, we sample three fixed variant-set sizes at *N* = 10, 50, and 100. We also test *N* = 500 across one draw for three Gemini models (Figure S1).

### Model inference and output parsing

We evaluate thirteen general-purpose language models: Claude Opus 5, GPT-5.6 Sol, GPT-5.5, GPT-5.4 mini, GPT-5.4 nano, Claude Opus 4.8, Claude Opus 4.7, Gemini 3.6 Flash, Gemini 3.5 Flash, Gemini 3.1 Pro, Gemini 3.1 Flash-Lite, GLM-5.2, and Kimi K3. Primary *N* = 50 runs use max reasoning for Sol, all three Opus models, and Kimi K3, xhigh for the other GPT models, and high for Gemini and GLM. All LLMs are evaluated with no external tools, internet access, or programmatic execution enabled. We also attempted to evaluate Fable 5, but Anthropic restricted all scoring attempts.

Models are instructed to return a JSON object of the form `{“ranking”: [“M03”, “M27”, …]}`. The parser looks exactly for a `“ranking”` array. A response is considered eligible if it contains at least 80% of the expected identifiers. Any omitted identifiers are appended after the explicit ranking in the prompt order. Every model evaluated on the primary *N* = 50 leaderboard contained 100% of all expected identifiers (Figure S3).

### Published-predictor evaluation

We compare against 95 predictors as baselines: 46 sequence-only methods and 49 methods that use alignments and/or structures. For each predictor, we subset scores from ProteinGym to match the same variant-sets in the benchmark. We take these scores and evaluate the resulting order with the same metric and aggregation used for the LLMs. Predictors use the inputs they were designed for, including, for some models, alignments and structures.

### Scoring and aggregation

We compare returned variant ranks with assay experimental values using Spearman rank correlation. A value of *ρ* = 1 indicates perfect ordering, whereas *ρ* = 0 indicates no monotonic rank association. This produces one score for each model, assay, and variant list. However, *ρ* = 1 is not a realistic estimate of the reproducibility ceiling of an assay. Experimental noise and differences between DMS studies mean that repeated measurements can disagree. ^21;22^

The primary leaderboard uses the published ProteinGym nested-macro aggregation. ^6^ We calculate the score separately for each variant list draw, then average the three draw-specific scores. Within a draw, we average repeated assays within each protein– functional-category cell, average proteins within each functional category, and give equal weight to the five category means. This prevents proteins with many assays or categories with many proteins from dominating the score.

## 3 Results

### Claude Opus 5 and GPT 5.6 Sol lead the LLM leaderboard but remain below specialist predictors

Claude Opus 5 leads the primary *N* = 50 leaderboard at *ρ* = 0.406 (SE across 3 draws, 0.002; Figure 1). GPT-5.6 Sol follows closely at 0.402. However, OpenAI and Anthropic refuse different assays. When both models are evaluated on the same assays, GPT 5.6 Sol leads at *ρ* = 0.409 versus *ρ* = 0.400 (Figure S2). In third place is Claude Opus 4.8 at 0.356, followed by Gemini 3.5 Flash at 0.343, GPT-5.5 at 0.336, and Gemini 3.1 Pro at 0.328 (Figure 1, Table S1).

**Figure 1.**
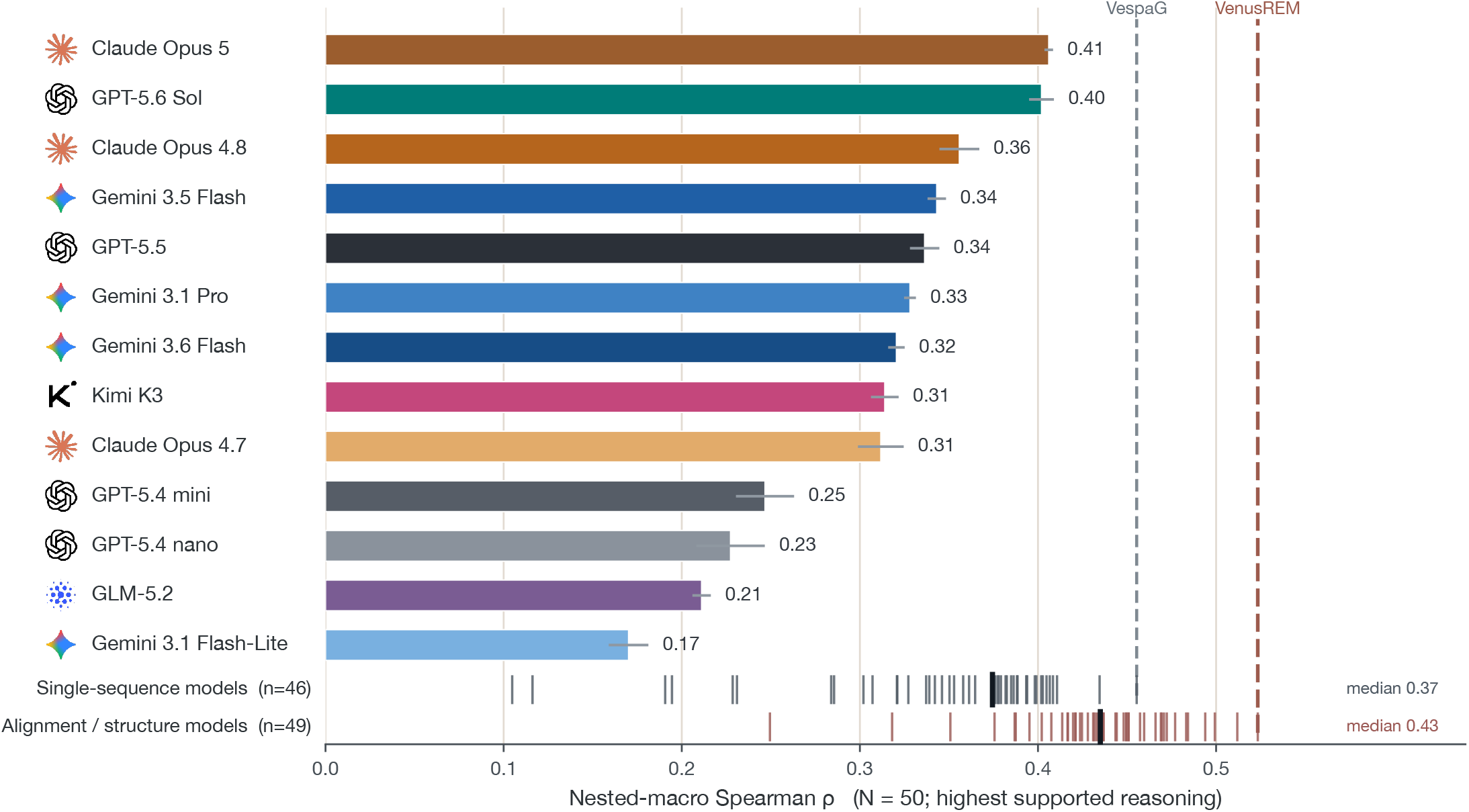
Leaderboard at N = 50. Nested-macro Spearman *ρ* for thirteen LLMs at their strongest reasoning settings; error bars show SE across three draws. Claude Opus 5 leads at 0.406, narrowly ahead of GPT-5.6 Sol at 0.402. The 95 rescored predictors are composed of 46 sequence-only methods (upper rug) and 49 methods using an alignment or structure (lower rug). Heavy ticks mark group medians (0.374 and 0.435); dashed lines mark VespaG (0.455) and VenusREM (0.523). ^1;2^

Opus 5 outperforms 49 of the 95 published comparators: 41 of 46 sequence-only methods and 8 of 49 methods using an alignment, structure, or both. Opus 5 and GPT 5.6 Sol beat the sequence-only median (*ρ* = 0.374) but remain below the alignment/structure median (*ρ* = 0.435). Opus 5 performs well below the best biomolecular model, VenusREM (*ρ* = 0.523). ^2^ If we combine the best scores from Opus 5 and VenusREM, this composite score improves on VenusREM by only 0.014, indicating limited complementarity (Figure S7).

Opus 5 and GPT-5.6 Sol score closely across taxonomic groups, but differ in their performance across functional categories (Figure S4, Figure S5). Opus 5 scores higher on activity (0.436 versus 0.422) and stability (0.484 versus 0.440), while Sol scores higher on binding (0.369 versus 0.316), expression (0.406 versus 0.393), and organismal fitness (0.375 versus 0.370). Stability is the strongest category for both models.

Increasing the number of variants makes the task harder for every LLM (Figure S1). The performance of GPT-5.6 Sol decreases from *ρ* = 0.423 at *N* = 10 to 0.375 at *N* = 100, while Gemini 3.1 Flash-Lite decreases from 0.300 to 0.083. In contrast, ESM2-150M, ESM2-650M, and VenusREM are nearly unchanged across variant-set sizes. This decline is consistent with limitations in current LLMs’ ability to effectively use large contexts when comparing a large number of variants.

### Test-time scaling improves performance but gains taper

Higher effort improves Spearman correlation in all four LLMs tested (Figure 2). GPT 5.6 Sol rises from 0.322 *±* 0.010 at low effort to 0.394 *±* 0.002 at xhigh and 0.402 *±* 0.007 at max. Opus 4.8 increases from 0.209 *±* 0.003 at low to 0.356 *±* 0.011 at max; GPT-5.5 from 0.248 *±* 0.013 at low to 0.336 *±* 0.008 at xhigh; and Gemini 3.5 Flash from 0.254 *±* 0.004 at low to 0.343 *±* 0.005 at high. Gains taper as reasoning effort increases. For further context, a difference of about 0.150 separates the top specialist from the 70th place specialist in ProteinGym.

**Figure 2.**
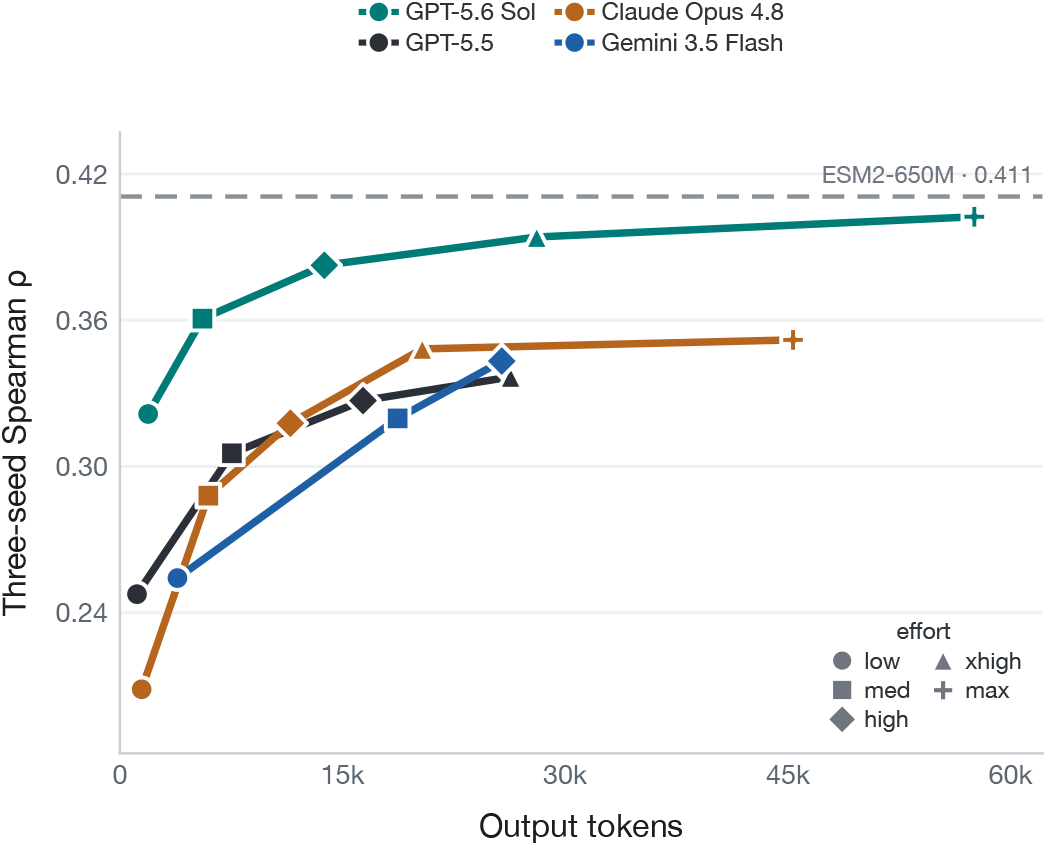
Test-time-compute scaling improves ranking before gains taper. N = 50 nested-macro *ρ* versus observed provider output tokens for Sol, GPT-5.5, Gemini 3.5 Flash, and Opus 4.8. The dashed line marks ESM2-650M on the primary N = 50 benchmark (*ρ* = 0.411). Marker shape identifies effort.

### Sequence-only methods score higher on high-depth assays

We stratify 180 assays based on alignment depth of each protein family. On average, LLM performance is unchanged between low- and high-depth proteins: 0.313 at low depth, 0.301 at medium depth, and 0.311 at high depth (Figure 3a). In contrast, the mean of 46 biomolecular sequence-only methods rises from 0.309 at low depth to 0.380 at medium depth and 0.411 at high depth (Figure 3a).

**Figure 3.**
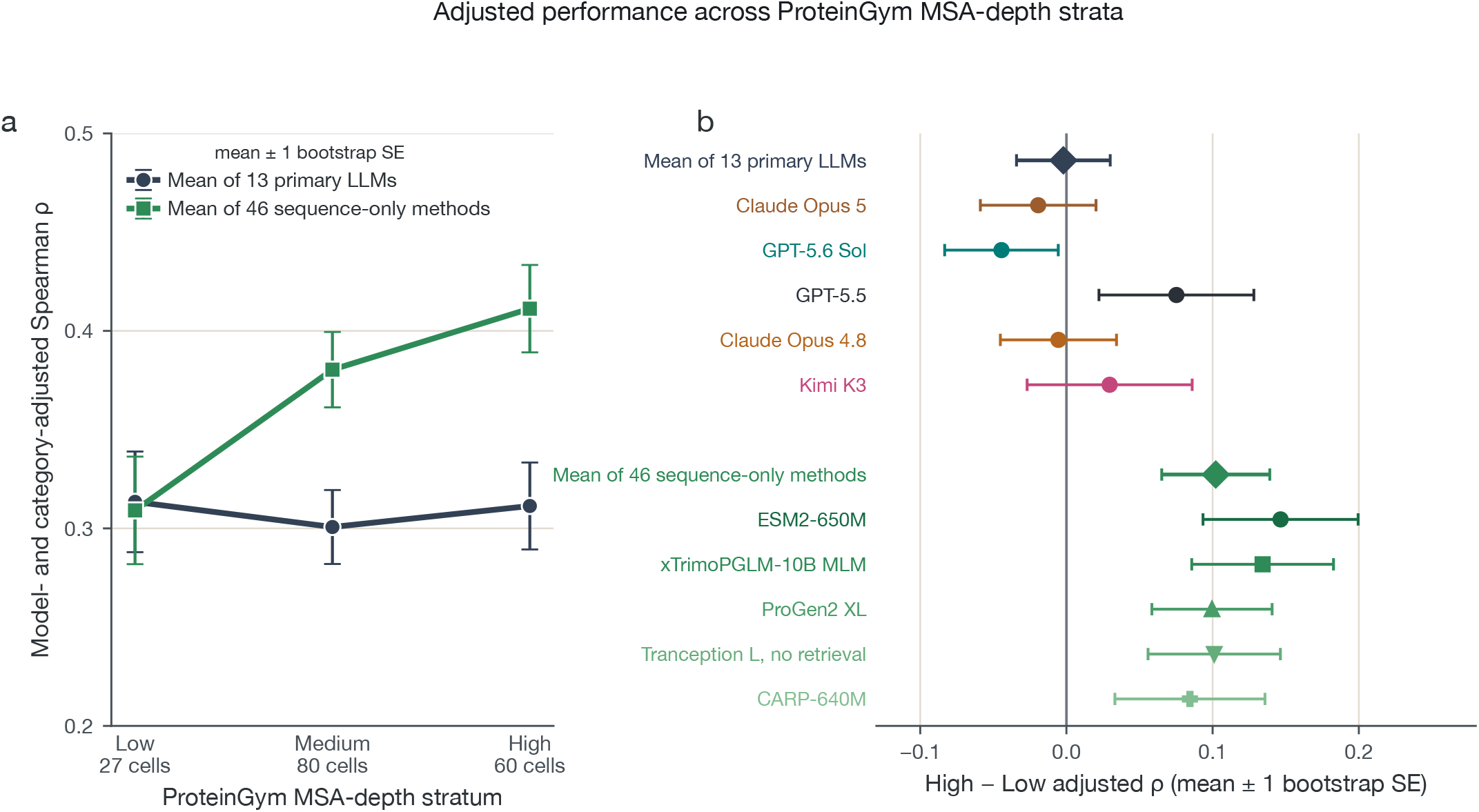
Sequence-only predictors improve with MSA depth, while LLMs do not. **(a)** Means for the thirteen-model primary LLM aggregate and the mean of 46 sequence-only methods. **(b)** High-minus-low contrasts for both aggregates, five representative LLMs (Opus 5, Sol, GPT-5.5, Opus 4.8, and Kimi K3), and representative sequence-only values from ESM2, xTrimoPGLM, ProGen2, retrieval-free Tranception, and CARP. ^12–15;23^ Error bars show *±*1 protein-cluster bootstrap SE.

This trend is also reproducible across individual LLMs and biomolecular models (Figure 3b). Sequence-only predictors perform better on proteins with deeper alignments, while LLM performance does not change with alignment depth.

### Source-study recognition is common but not positively associated with accuracy

ProteinGym is public, so models may recognize its source studies. To test the impact of recognition on performance, we audit all leaderboard traces for recognition with Gemini 3.5 Flash as an LLM judge. ^24^ Traces are marked positively if they identify the source study, dataset, or claim to recall a measured variant result.

Recognition is common but varies by model. It appears in 88% of Sol traces (512/585), 57% of Opus 4.8 traces (324/568), 45% of GPT-5.5 traces (268/595), and 9% of Gemini 3.5 Flash traces (61/651; Figure 4a). Most positive traces name a source study rather than claim recognition of a specific measurement. For example, Opus 5 identifies the NusA stability task as a Tsuboyama et al. 2023 dataset.

**Figure 4.**
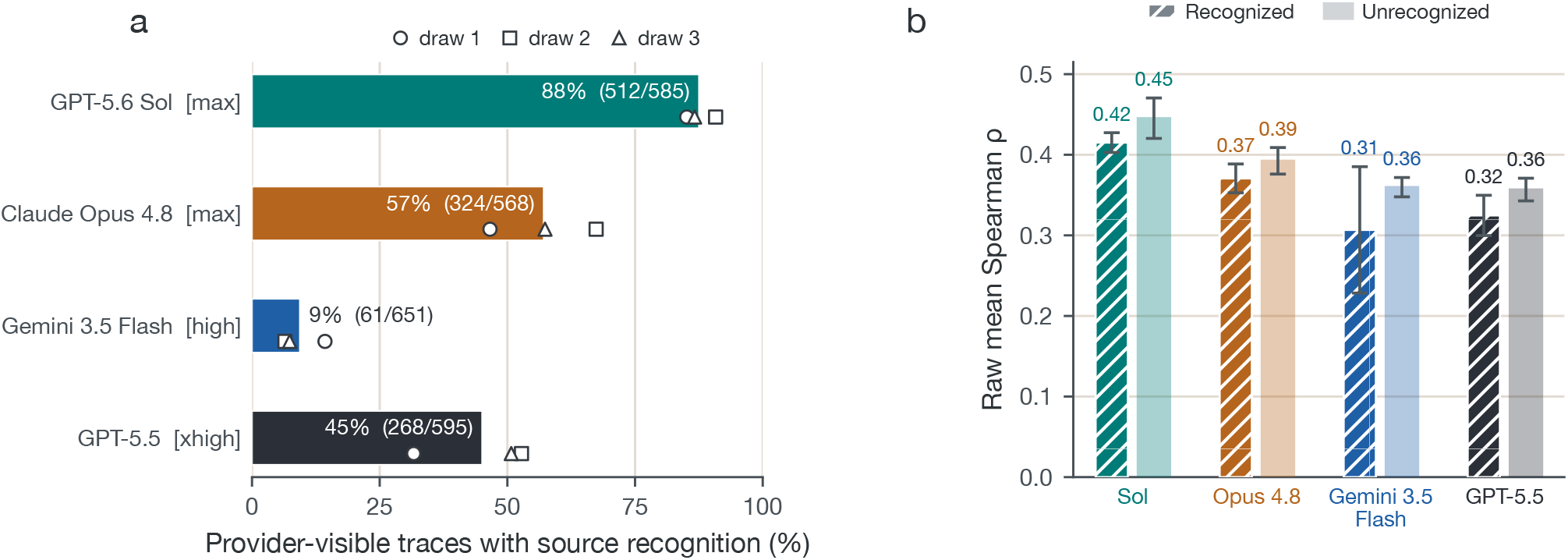
Study recognition is common but not positively associated with accuracy. **(a)** Bars show the average of recognition rates over every extractable trace; white markers show the three draws and labels give recognized/all traces. **(b)** Mean *ρ* for recognized traces (hatched) and unrecognized traces (lighter bars). Error bars show *±*1 protein-cluster bootstrap SE.

**Figure 5.**
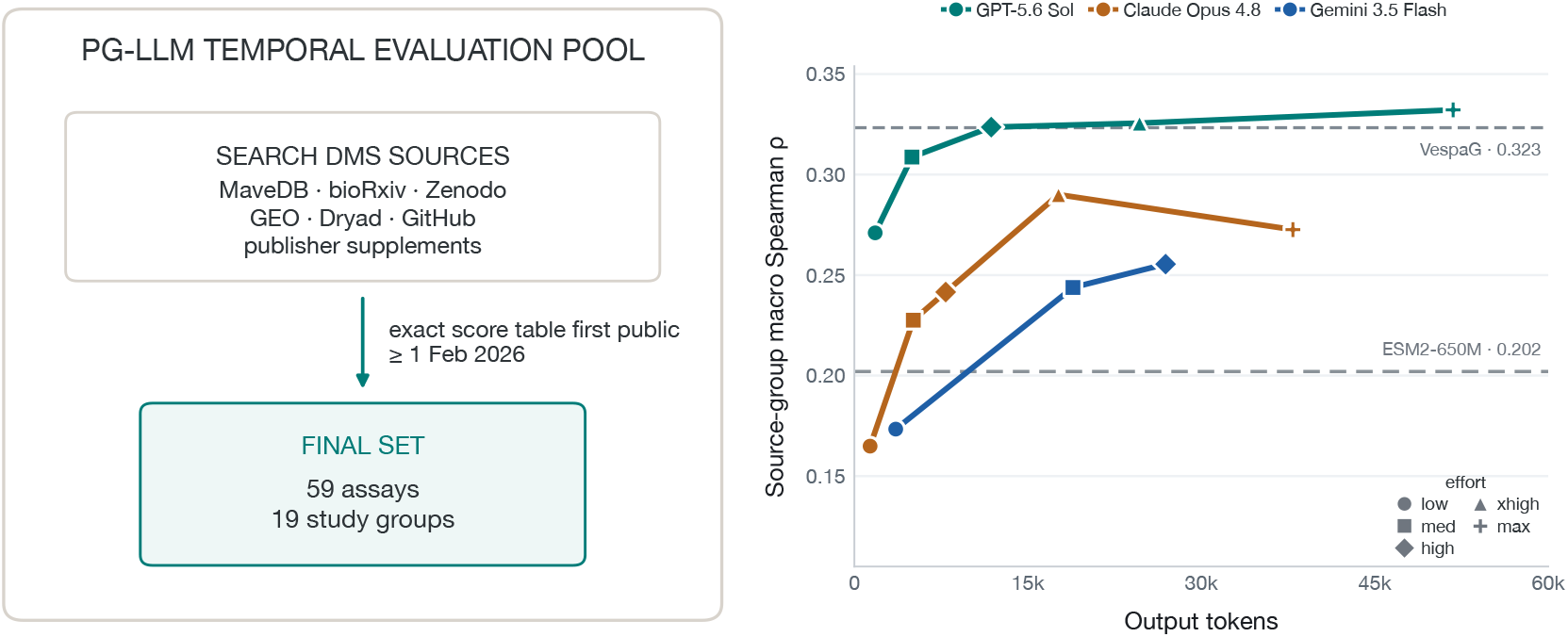
A post-knowledge-cutoff set of DMS assays shows similar LLM performance trends. **(a)** We searched MaveDB, bioRxiv, Zenodo, GEO, Dryad, GitHub, and publisher supplements for exact variant-level score tables first made public on or after 1 February 2026. **(b)** *N* = 50 Spearman *ρ* versus output tokens; points show means across three draws. ESM2-650M and VespaG v2 are shown as reference lines.

For each model and cell, we subtract the mean score of the other three models from that model’s score. This controls for the difficulty of the variant list being evaluated. We then take the residual scores and compare them between recognized and unrecognized traces. The differences are +0.011 for Sol, −0.003 for Opus 4.8, −0.016 for Gemini 3.5 Flash, and −0.051 for GPT-5.5, indicating no consistent positive association between recognition and accuracy. We also observe differences in the strategies described by each model, including their use of structural hypotheses, numeric scoring, and stabilizing-mutation heuristics (Figure S6).

### A post-knowledge-cutoff DMS assay set shows similar trends in performance

To better control for contamination risk, we construct a post-knowledge-cutoff evaluation set of DMS assays where the exact variant-level score tables became publicly retrievable on or after 1 February 2026. We identify 59 assays from 19 studies that meet this cutoff by searching MaveDB, bioRxiv, Zenodo, GEO, Dryad, GitHub, and publisher supplements (Table S4).

We evaluate GPT-5.6 Sol, Gemini 3.5 Flash, and Claude Opus 4.8 using the same *N* = 50, no-tools task, with a sweep across reasoning efforts. Studies included are after the knowledge cutoff of Gemini 3.5 Flash and Opus 4.8. ^25;26^ 57 of 59 assays also postdate Sol’s 16 February 2026 cutoff. ^27**?**^. Across all evaluated traces, we observe zero instances of study identification.

To prevent results from the same study from dominating the evaluation, we first average scores within each study-group and draw, then average across study-groups. On this set of 59 assays, Sol rises from 0.271 at low effort to 0.332 at max. Opus 4.8 rises from 0.165 at low to 0.290 at xhigh, then reaches 0.273 at max. Gemini 3.5 Flash rises from 0.173 at low to 0.255 at high. Despite lower absolute scores compared to ProteinGym assays, LLMs perform better relative to biomolecular comparators, with GPT 5.6 Sol (Max) scoring better than ESM2-650M (0.202) and being approximately on par with VespaG (0.323).

## 4 Discussion

PG-LLM shows that frontier LLMs place near the median of specialized protein predictors. Claude Opus 5 and GPT 5.6 Sol perform the best of all LLMs evaluated, with Opus 5 outperforming 49 of the 95 published comparators despite receiving only an assay description and full-length sequences. Current LLMs can reason well about a small set of variants, but reasoning becomes harder as the number of comparisons grows, consistent with evidence that LLMs use long contexts unevenly. ^28^ As models become more capable at the *N* = 50 setting, we plan to scale the benchmark standard to *N* = 100, then *N* = 500, and eventually to ranking every measured variant in each assay.

The MSA-depth analysis shows that sequence-only protein models improve on high-depth families, as expected if evolutionary sampling supplies useful information. ^7;11^ LLMs show no corresponding trend. General-purpose models may instead rely more on knowledge acquired from their broad pretraining corpora, such as the generic effects of charge, packing, and polarity.

Tokenization may also contribute to this gap. The protein language models tested here represent sequences one amino acid at a time. General-purpose LLMs use tokenizers designed for text, which likely divide protein sequences into multi-residue fragments. For an LLM, a single mutation can change several token boundaries, adding a representation problem before the model can reason about its biological effect. We do not isolate for this effect here, but it may help explain why general-purpose LLMs struggle to compare closely related sequences.

Source recognition in LLM traces does not show a consistent positive association with ranking accuracy. Models frequently name source studies or datasets, yet recognized traces are not more accurate after controlling for variant-set difficulty. However, this does not rule out training exposure or silent recall, and visible reasoning should not be treated as a fully faithful account of model reasoning. ^29;30^

The recent-assay analysis helps mitigate concerns about contamination in the results presented here. In our post-knowledge-cutoff set, we observe similar test-time compute scaling across all three model families, and GPT 5.6 Sol at max performs approximately on par with VespaG.

## 5 Limitations

PG-LLM evaluates controlled variant sets rather than complete assays. The primary benchmark uses three samples of 50 variants per assay, while the standard ProteinGym benchmark scores each method over the full measured substitution set. Therefore, absolute values of scores discussed here are not interchangeable with ProteinGym scores.

LLMs are only given natural-language context and do not receive alignments or structures. Future experiments should test the addition of alignments, structures, or protein-analysis tools to test whether language models can integrate the evidence that currently gives specialist systems their advantage.

ProteinGym may have appeared in model training. We make use of opaque shuffled labels to make direct answer reuse difficult, but this cannot fully prevent recognition of a protein, study, or measured variant. The trace audit we conduct only captures explicit information in provider-visible text and cannot establish training membership or exclude silent recall.

Provider safety policies and refusals change model coverage. Opus 5 leads on each model’s raw scored assays, while Sol leads on both the shared 180-assay set between the two models.

PG-LLM covers ranking of measured substitution variants; it does not evaluate indels, clinical pathogenicity, wet-lab hit rate, or learning across an iterative optimization campaign.

## 6 Resources and reproducibility

Aggregate results, assay-level predictions, and traces are available at ProteinGymLLM.com. Code to run the PG-LLM benchmark is released through the PG-LLM GitHub repository.

## Acknowledgements

We thank Dr. Anthony Gitter for feedback on the manuscript and blog post. We thank OpenAI for compute credits, Modal for compute credits, and otto-SR for supporting this work. Rohit Arora is supported by the Canadian Institutes of Health Research (CIHR). The funders had no role in prompt design or result reporting.

## Supplementary

**Table S1.** Primary N = 50 results by draw. Each draw column reports nested-macro Spearman *ρ* for one fixed variant-set. The mean averages the three draw-specific nested-macro scores. Assays is the number of datasets with at least one scored draw; highlighting marks the maximum in each score column.

| Model | Effort | Draw 1 $\rho$ | Draw 2 $\rho$ | Draw 3 $\rho$ | Mean $\rho$ | Assays |
| --- | --- | --- | --- | --- | --- | --- |
| Claude Opus 5 | max | **0.402** | **0.405** | 0.411 | **0.406** | 188 |
| GPT-5.6 Sol | max | 0.393 | 0.397 | **0.416** | 0.402 | 196 |
| Claude Opus 4.8 | max | 0.333 | 0.368 | 0.366 | 0.356 | 185 |
| Gemini 3.5 Flash | high | 0.342 | 0.335 | 0.353 | 0.343 | 217 |
| GPT-5.5 | xhigh | 0.334 | 0.324 | 0.352 | 0.336 | 200 |
| Gemini 3.1 Pro | high | 0.331 | 0.322 | 0.332 | 0.328 | 217 |
| Gemini 3.6 Flash | high | 0.327 | 0.312 | 0.323 | 0.321 | 217 |
| Kimi K3 | max | 0.302 | 0.312 | 0.328 | 0.314 | 217 |
| Claude Opus 4.7 | max | 0.296 | 0.302 | 0.337 | 0.312 | 184 |
| GPT-5.4 mini | xhigh | 0.235 | 0.226 | 0.279 | 0.247 | 201 |
| GPT-5.4 nano | xhigh | 0.230 | 0.193 | 0.259 | 0.227 | 217 |
| GLM-5.2 | high | 0.204 | 0.221 | 0.208 | 0.211 | 217 |
| Gemini 3.1 Flash-Lite | high | 0.178 | 0.148 | 0.184 | 0.170 | 217 |

**Figure S1.**
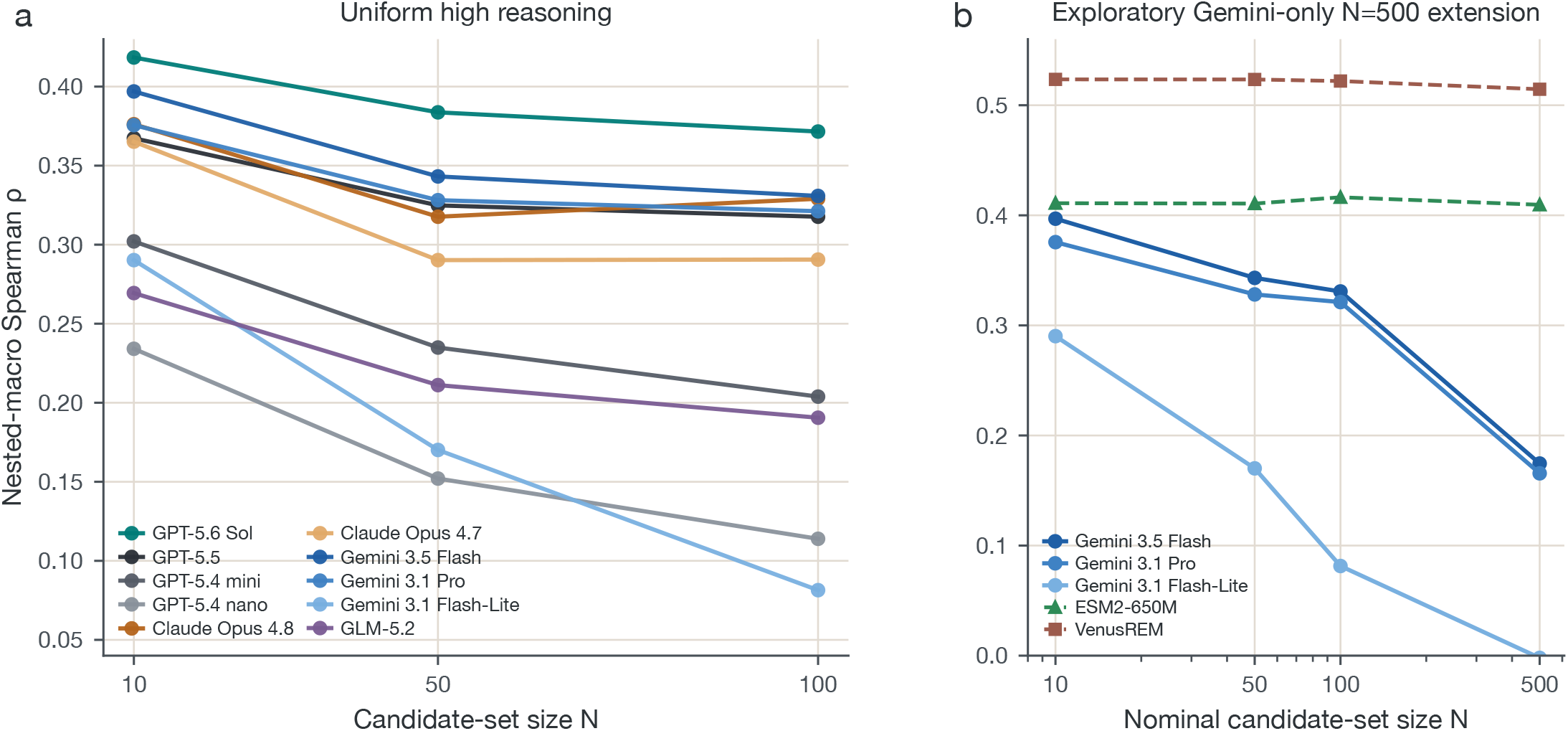
Variant-set sensitivity at uniform high reasoning. **(a)** Nested-macro *ρ* at N = 10, 50, and 100 for ten models. **(b)** Gemini-only extension to N = 500. N = 500 uses one draw and includes assays with fewer than 500 available variants.

**Figure S2.**
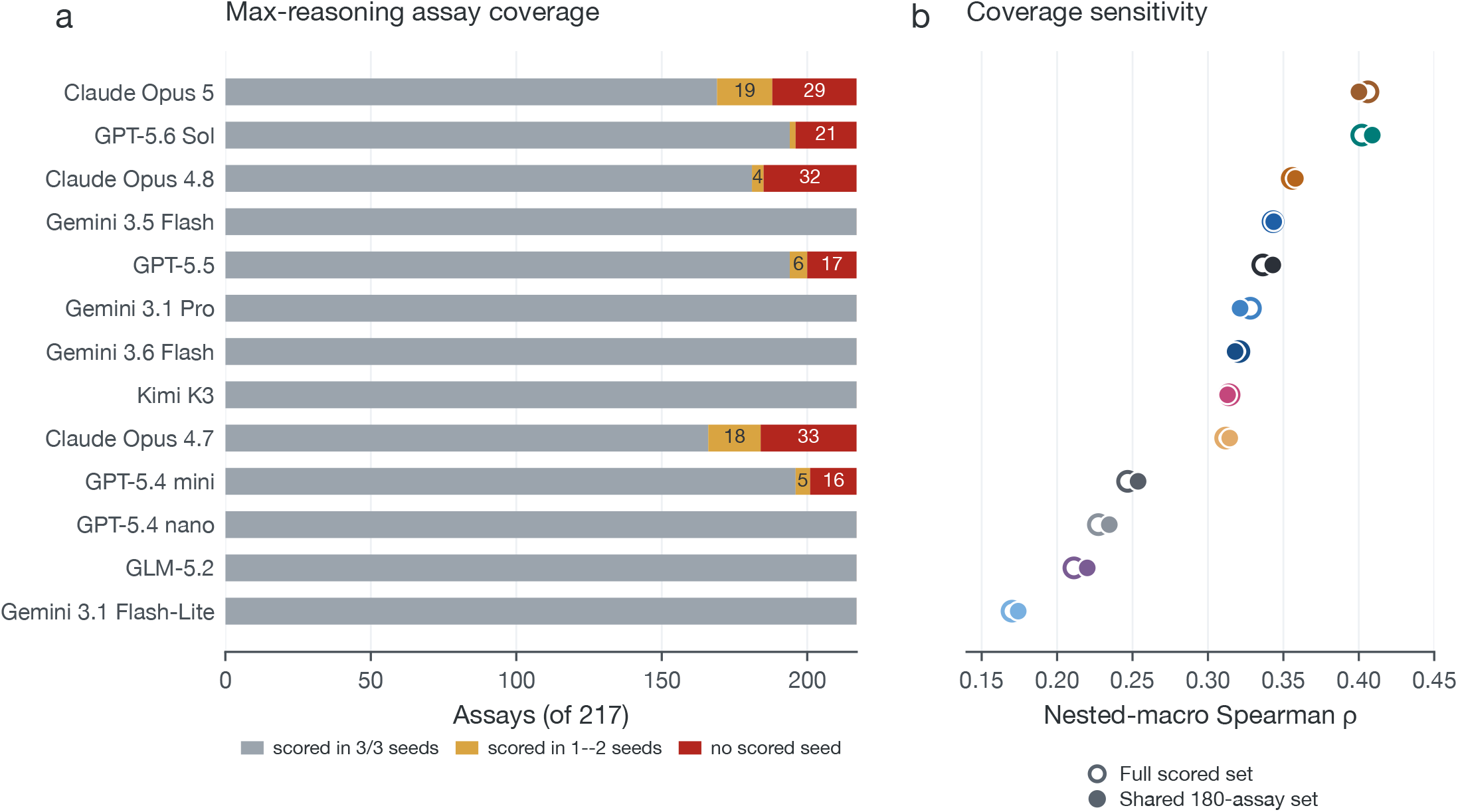
Provider outcomes make the top-two ordering coverage-sensitive. **(a)** Gray bars mark assays scored in all three draws; gold and red segments show partial or absent coverage among 217 assays. **(b)** Model-colored open circles show each model’s full scored set; closed circles show the shared 180-assay set. The broad ordering of the remaining models is largely stable.

**Figure S3.**
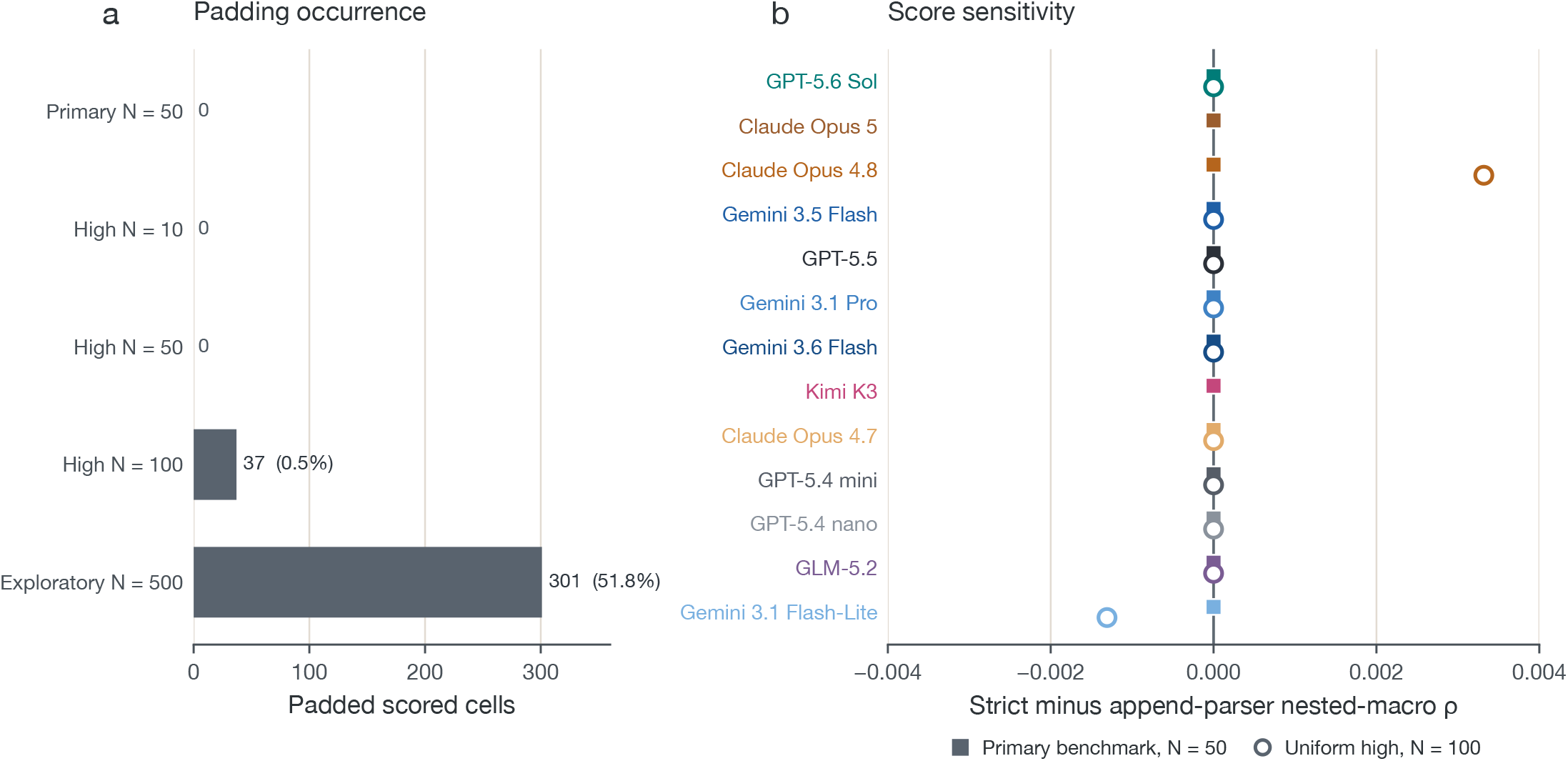
Lack of parser completion is rare and has little effect at N = 100. **(a)** Padded scored cells in each variant-set size. The N = 50 contains no padded cells. Uniform-high N = 100 contains 37 of 6,790 (0.5%), whereas exploratory N = 500 contains 301 of 581 (51.8%). **(b)** Strict-minus-append-parser score shifts. Squares show N = 50 models. Open circles show the eleven-model uniform-high N = 100 sensitivity

**Figure S4.**
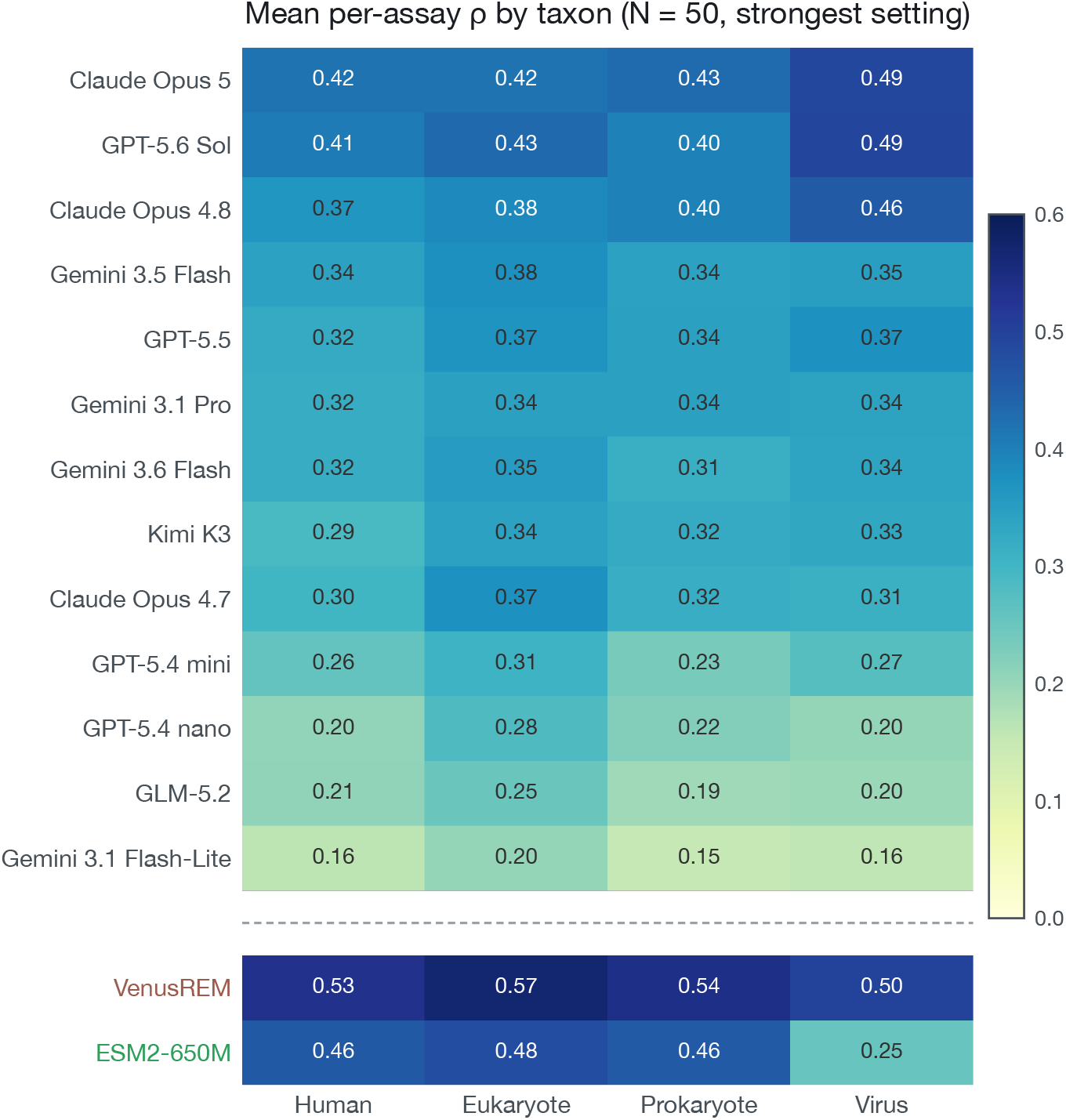
Leaderboard accuracy by taxonomic group. Mean N = 50 Spearman *ρ* within four ProteinGym taxonomic groups at each LLM’s strongest supported setting; VenusREM and ESM2-650M appear below the divider.

**Figure S5.**
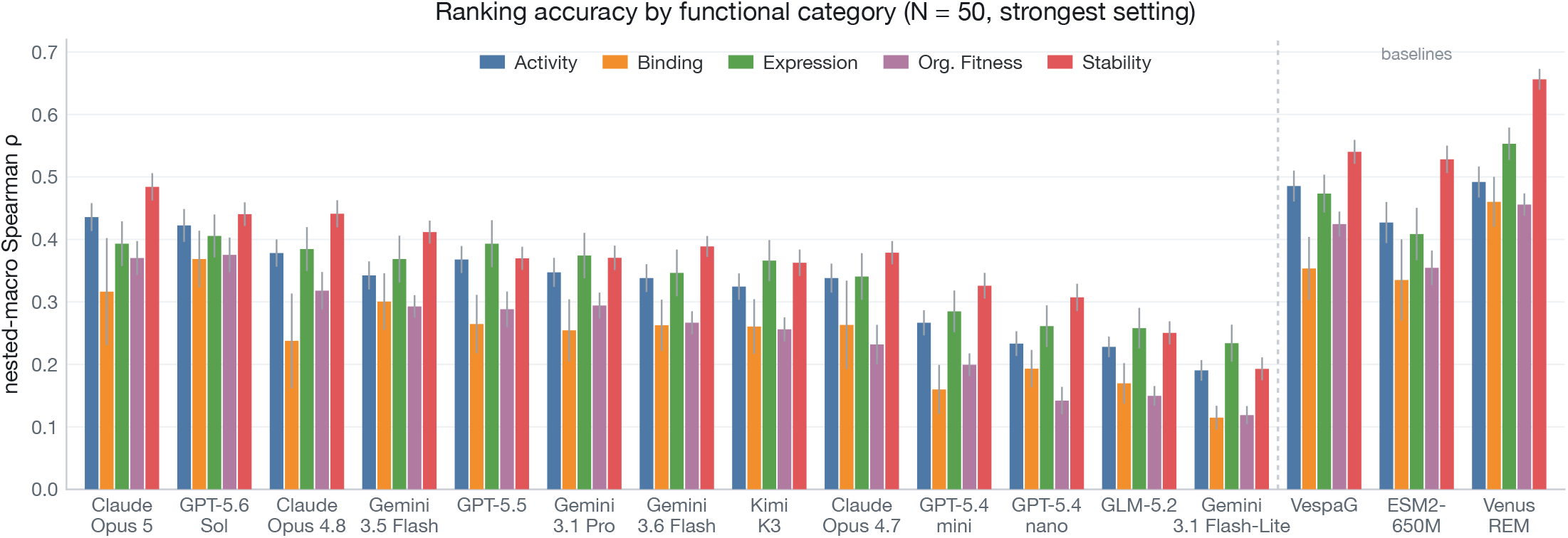
Primary accuracy by functional category. N = 50 Spearman *ρ* within each ProteinGym functional category at every LLM’s strongest supported setting; VespaG, ESM2-650M, and VenusREM appear to the right of the divider. Thin lines show protein-cluster bootstrap SE.

**Figure S6.**
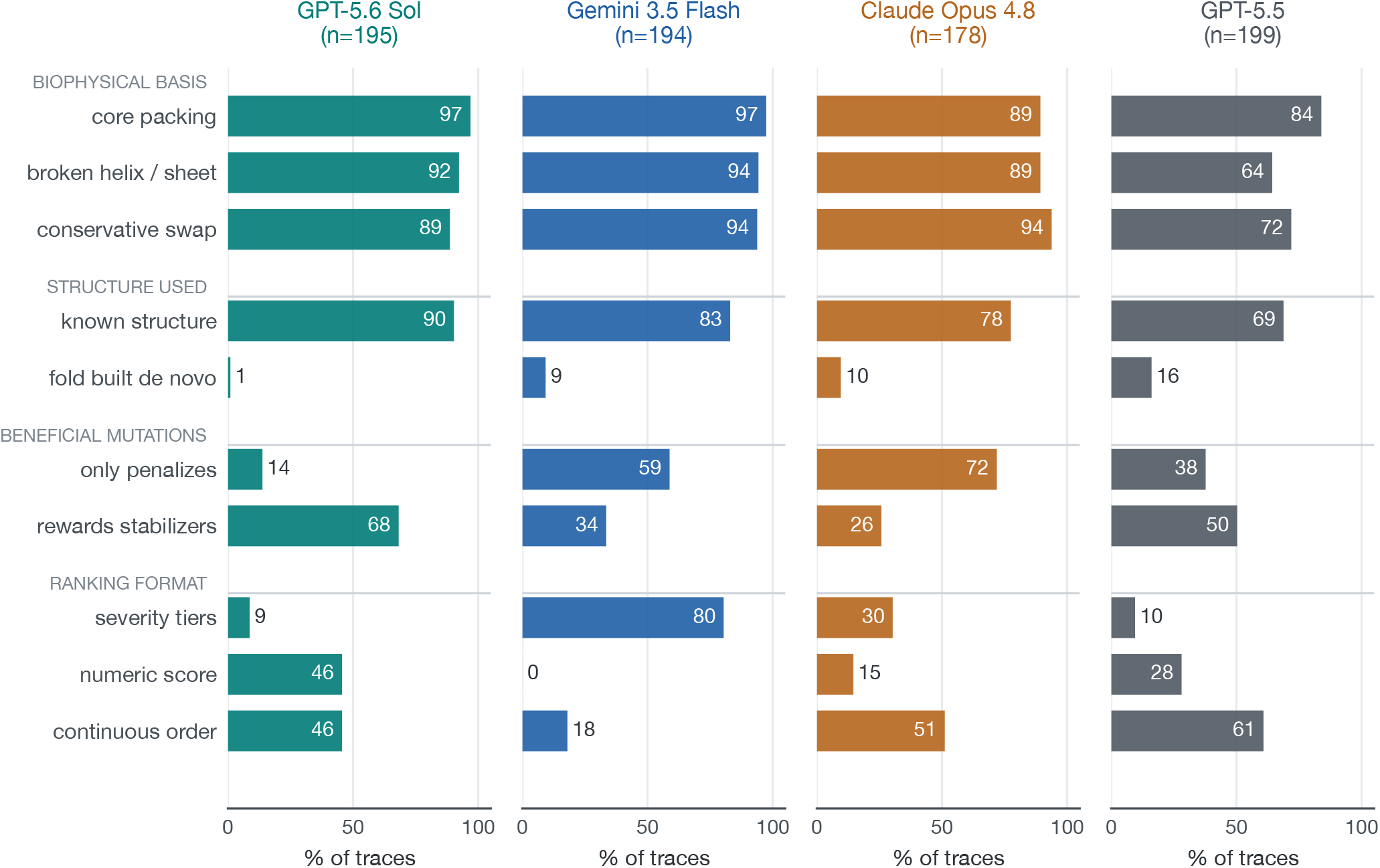
Rationale-content analysis. Bars show the percentage of visible traces assigned each rubric code on a common 0–100% scale. These frequencies describe visible text.

**Figure S7.**
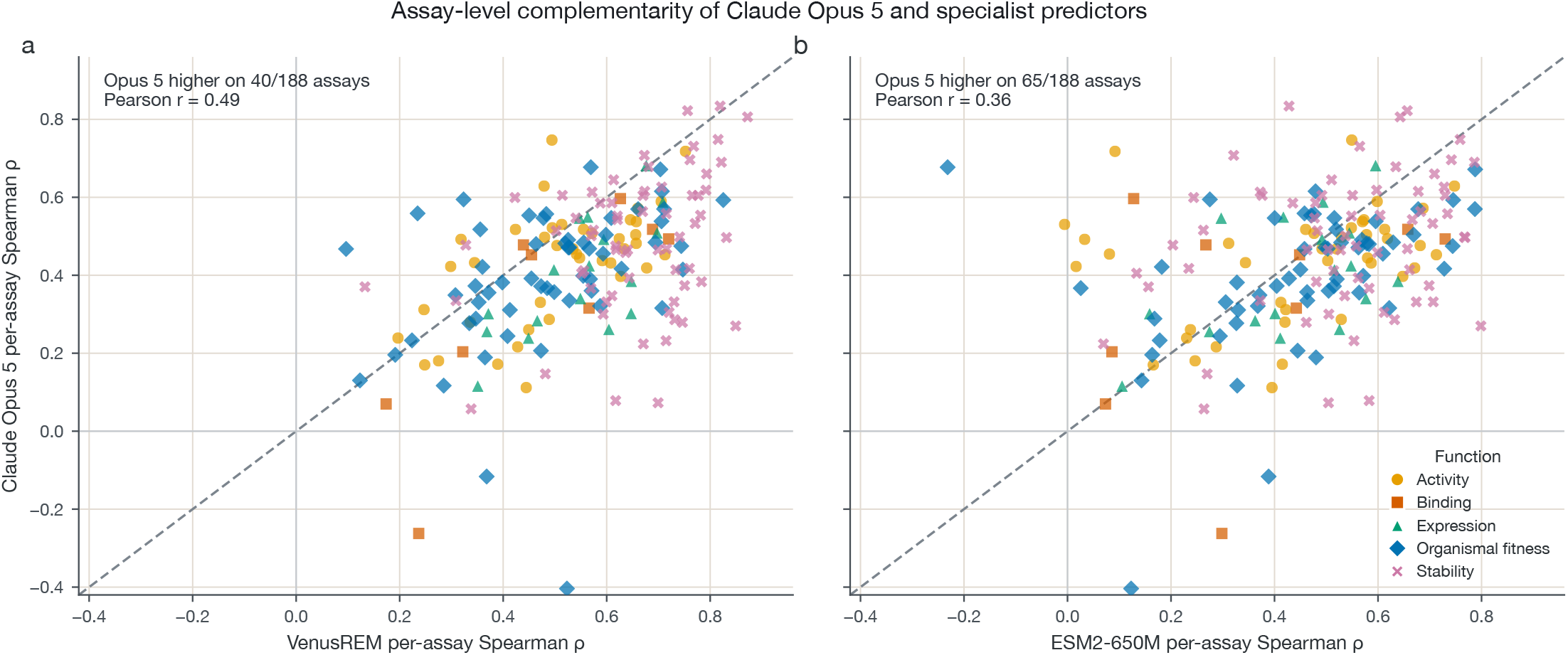
Opus 5 wins on some assays, but an oracle assay router adds little. Each point is one N = 50 assay, colored by functional category. Max-reasoning Opus 5 is compared with **(a)** VenusREM and **(b)** ESM2-650M. Across 188 shared assays, Opus 5 is higher on 40 against VenusREM (*r* = 0.49) and 65 against ESM2-650M (*r* = 0.36).

**Table S2.** High-effort results at N = 10 and N = 100. Score is Spearman *ρ*; SE is the standard error across three fixed variant draws. Highlighting marks the maximum in each score column.

| (a) N = 10 |  |  |  |  |  |  |  |  |  |
| --- | --- | --- | --- | --- | --- | --- | --- | --- | --- |
| Model | Score | SE | Single | Multi | Activity | Binding | Expression | Org. fit. | Stability |
| GPT-5.6 Sol | **0.418** | 0.017 | 0.395 | **0.533** | 0.418 | **0.420** | 0.426 | **0.328** | **0.500** |
| Gemini 3.5 Flash | 0.397 | 0.008 | **0.397** | 0.370 | **0.444** | 0.343 | **0.448** | 0.276 | 0.474 |
| Claude Opus 4.8 | 0.382 | 0.015 | 0.347 | 0.446 | 0.402 | 0.319 | 0.442 | 0.282 | 0.466 |
| Gemini 3.1 Pro | 0.376 | 0.022 | 0.380 | 0.353 | 0.363 | 0.358 | 0.420 | 0.315 | 0.422 |
| GPT-5.5 | 0.367 | 0.000 | 0.370 | 0.370 | 0.389 | 0.296 | 0.422 | 0.288 | 0.442 |
| Claude Opus 4.7 | 0.363 | 0.014 | 0.359 | 0.333 | 0.384 | 0.358 | 0.388 | 0.245 | 0.442 |
| GPT-5.4 mini | 0.303 | 0.023 | 0.289 | 0.303 | 0.338 | 0.221 | 0.352 | 0.275 | 0.328 |
| Gemini 3.1 Flash-Lite | 0.290 | 0.013 | 0.276 | 0.266 | 0.322 | 0.251 | 0.311 | 0.210 | 0.358 |
| GLM-5.2 | 0.269 | 0.019 | 0.264 | 0.250 | 0.300 | 0.208 | 0.287 | 0.221 | 0.331 |
| GPT-5.4 nano | 0.234 | 0.015 | 0.212 | 0.270 | 0.245 | 0.189 | 0.309 | 0.161 | 0.267 |

| (b) N = 100 |  |  |  |  |  |  |  |  |  |
| --- | --- | --- | --- | --- | --- | --- | --- | --- | --- |
| Model | Score | SE | Single | Multi | Activity | Binding | Expression | Org. fit. | Stability |
| GPT-5.6 Sol | **0.371** | 0.001 | **0.364** | **0.430** | **0.397** | **0.343** | **0.402** | **0.335** | 0.381 |
| Gemini 3.5 Flash | 0.331 | 0.005 | 0.324 | 0.308 | 0.346 | 0.292 | 0.347 | 0.284 | 0.385 |
| Claude Opus 4.8 | 0.330 | 0.006 | 0.326 | 0.337 | 0.358 | 0.290 | 0.343 | 0.268 | **0.390** |
| Gemini 3.1 Pro | 0.321 | 0.001 | 0.321 | 0.317 | 0.336 | 0.275 | 0.351 | 0.286 | 0.357 |
| GPT-5.5 | 0.321 | 0.004 | 0.323 | 0.325 | 0.359 | 0.259 | 0.380 | 0.281 | 0.327 |
| Gemini 3.6 Flash | 0.319 | 0.007 | 0.299 | 0.388 | 0.339 | 0.268 | 0.330 | 0.269 | 0.387 |
| Claude Opus 4.7 | 0.289 | 0.002 | 0.272 | 0.280 | 0.304 | 0.278 | 0.299 | 0.233 | 0.333 |
| GPT-5.4 mini | 0.205 | 0.004 | 0.187 | 0.207 | 0.209 | 0.145 | 0.216 | 0.184 | 0.270 |
| Gemini 3.1 Flash-Lite | 0.081 | 0.013 | 0.082 | 0.062 | 0.094 | 0.092 | 0.089 | 0.055 | 0.077 |
| GLM-5.2 | 0.191 | 0.009 | 0.168 | 0.181 | 0.197 | 0.178 | 0.211 | 0.148 | 0.218 |
| GPT-5.4 nano | 0.114 | 0.024 | 0.088 | 0.123 | 0.107 | 0.071 | 0.136 | 0.062 | 0.193 |

**Table S3.** Primary N = 50 cell outcomes. Counts cover 217 assays and three draws (651 expected cells per model). Overflow is a request that remained truncated at its supported ceiling.

| Model | Scored | Safety block | Refusal | Overflow | Other |
| --- | --- | --- | --- | --- | --- |
| Claude Opus 5 | 541 | 0 | 110 | 0 | 0 |
| GPT-5.6 Sol | 585 | 60 | 0 | 0 | 6 |
| GPT-5.5 | 591 | 54 | 0 | 0 | 6 |
| GPT-5.4 mini | 595 | 50 | 0 | 0 | 6 |
| GPT-5.4 nano | 651 | 0 | 0 | 0 | 0 |
| Claude Opus 4.8 | 550 | 0 | 101 | 0 | 0 |
| Claude Opus 4.7 | 525 | 0 | 125 | 1 | 0 |
| Gemini 3.5 Flash | 651 | 0 | 0 | 0 | 0 |
| Gemini 3.1 Pro | 651 | 0 | 0 | 0 | 0 |
| Gemini 3.6 Flash | 651 | 0 | 0 | 0 | 0 |
| Kimi K3 | 651 | 0 | 0 | 0 | 0 |
| Gemini 3.1 Flash-Lite | 651 | 0 | 0 | 0 | 0 |
| GLM-5.2 | 651 | 0 | 0 | 0 | 0 |

**Table S4.** Sources in the recent-assay temporal panel. The panel contains 59 assays from 19 deduplicated study or data-source groups; first-public date is the earliest audited public release of the exact variant-level score table.

| First-public date | Study title | *n* | Reference |
| --- | --- | --- | --- |
| 2026-02-09 | Comprehensive Variant Effect Map of Parkin-Mediated Mitophagy in Parkinson's Disease | 1 | 31 |
| 2026-02-09 | A Functional Genetic Atlas of Parkin Resolves Variants of Uncertain Significance and Predicts Parkinson's Disease Age at Onset | 1 | 32 |
| 2026-02-19 | Systematic mutational mapping reveals optimal amyloid formation for RIPK function | 4 | 33 |
| 2026-03-30 | Determinants of metal import and specificity in a bacterial transporter | 1 | 34 |
| 2026-04-12 | Mutational Scanning of α-Synuclein using a Clickable Protein Tag Reveals Determinants of Membrane-Induced Aggregation | 1 | 35 |
| 2026-04-21 | Next generation sequence counts for different *E. coli* Cra mutant libraries after cell survival selection | 1 | 36 |
| 2026-04-23 | Landscape-scale navigation unlocks antibody CDR structural logic for AI-guided rescue and therapeutic optimization | 17 | 37 |
| 2026-05-06 | Interactions of outer membrane lipoproteins *P. aeruginosa* PA3214 and *E. coli* PqiC with their MCE protein binding partners, PA3213 and PqiB | 1 | 38 |
| 2026-05-07 | GROQ-seq Function Measurements for LacI, RamR, and VanR Transcription Factor Libraries | 5 | 39 |
| 2026-05-11 | Redesign of energetically frustrated regions rescues function in defective T4 clamp loaders | 1 | 40 |
| 2026-05-12 | GROQ-seq Function Measurements for TEV Protease SSVL and epPCR Libraries | 1 | 41 |
| 2026-06-11 | Multimodal phenotyping defines variant-to-function maps for RBM20 in dilated cardiomyopathy | 2 | 42 |
| 2026-06-16 | Structural and Functional Principles of Hcp-Mediated Antibacterial Toxin Delivery by the Type VI Secretion System | 1 | 43 |
| 2026-06-24 | The mutational landscape of STING-induced immunity | 2 | 44 |
| 2026-07-01 | Environment-dependent landscapes of coding variant impacts on coproporphyrinogen oxidase | 2 | 45 |
| 2026-07-03 | Protein Surface Site Determines the Evolutionary Accessibility of Allosteric Regulation | 1 | 46 |
| 2026-07-06 | An Atlas of Short Linear Motif-Mediated Human Protein-Protein Interactions | 15 | 47 |
| 2026-07-08 | Machine learning guided cell-free expression maps the biochemical landscape of carbonic anhydrase | 1 | 48 |
| 2026-07-10 | Genotype-phenotype single-cell transcriptomics for massive parallel assessment of genetic variants | 1 | 49 |

## References

1. Céline Marquet, Julius Schlensok, Marina Abakarova, Burkhard Rost, and Elodie Laine. Expert-guided protein language models enable accurate and blazingly fast fitness prediction. Bioinformatics, 40(11):btae621, 11 2024. ISSN 1367-4811. doi: 10.1093/bioinformatics/btae621. URL https://doi.org/10.1093/bioinformatics/btae621.

2. Yang Tan, Ruilin Wang, Banghao Wu, Liang Hong, and Bingxin Zhou. From high-throughput evaluation to wet-lab studies: advancing mutation effect prediction with a retrieval-enhanced model. Bioinformatics, 41(Supplement 1):i401–i409, 07 2025. ISSN 1367-4811. doi: 10.1093/bioinformatics/btaf189. URL https://doi.org/10.1093/bioinformatics/btaf189.

3. Aashka Bhowmick, Nasim Rahmatpour, Shivang Sharma, Qian Xu, Ananya Gupta, Asmita Lagwankar, Arjun Banerjee, and Christopher Zou. VariantBench: An agentic benchmark for genetic variant discovery and interpretation, 2026. URL https://github.com/latchbio/variantbench.

4. Jeremy Li, Suyash Shringarpure, Edmund Wong, and Andrew Ho. GeneBench-Pro: Evaluating Multistage Statistical Reasoning in Genomics, Quantitative Biology, and Translational Biomedicine. bioRxiv, 2026. doi: 10.64898/2026.06.29.735386. URL https://www.biorxiv.org/content/early/2026/07/21/2026.06.29.735386.

5. D. M. Fowler and S. Fields. Deep mutational scanning. Nat. Methods, 11:801–807, 2014.

6. Pascal Notin, Aaron Kollasch, Daniel Ritter, Lood van Niekerk, Steffanie Paul, Han Spinner, Nathan Rollins, Ada Shaw, Rose Orenbuch, Ruben Weitzman, Jonathan Frazer, Mafalda Dias, Dinko Franceschi, Yarin Gal, and Debora Marks. ProteinGym: Large-scale benchmarks for protein fitness prediction and design. In A. Oh, T. Neumann, A. Globerson, K. Saenko, M. Hardt, and S. Levine, editors, Advances in Neural Information Processing Systems, volume 36, pages 64331–64379. Curran Associates, Inc., 2023. URL https://proceedings.neurips.cc/paper_files/paper/2023/file/cac723e5ff29f65e3fcbb0739ae91bee-Paper-Datasets_and_Benchmarks.pdf.

7. Thomas A Hopf, John B Ingraham, Frank J Poelwijk, Charlotta P I Schärfe, Michael Springer, Chris Sander, and Debora S Marks. Mutation effects predicted from sequence co-variation. Nature Biotechnology, 35(2):128–135, February 2017. ISSN 1546-1696. doi: 10.1038/nbt.3769. URL https://doi.org/10.1038/nbt.3769.

8. Adam J. Riesselman, John B. Ingraham, and Debora S. Marks. Deep generative models of genetic variation capture the effects of mutations. Nature Methods, 15 (10):816–822, October 2018. ISSN 1548-7105. doi: 10.1038/s41592-018-0138-4. URL https://doi.org/10.1038/s41592-018-0138-4.

9. Jonathan Frazer, Pascal Notin, Mafalda Dias, Aidan Gomez, Joseph K. Min, Kelly Brock, Yarin Gal, and Debora S. Marks. Disease variant prediction with deep generative models of evolutionary data. Nature, 599(7883):91–95, November 2021. ISSN 1476-4687. doi: 10.1038/s41586-021-04043-8. URL https://doi.org/10.1038/s41586-021-04043-8.

10. E. Laine, Y. Karami, and A. Carbone. GEMME: A simple and fast global epistatic model predicting mutational effects. Mol. Biol. Evol., 36:2604–2619, 2019.

11. Roshan M Rao, Jason Liu, Robert Verkuil, Joshua Meier, John Canny, Pieter Abbeel, Tom Sercu, and Alexander Rives. MSA transformer. In Marina Meila and Tong Zhang, editors, Proceedings of the 38th International Conference on Machine Learning, volume 139 of Proceedings of Machine Learning Research, pages 8844–8856. PMLR, 18–24 Jul 2021. URL https://proceedings.mlr.press/v139/rao21a.html.

12. Zeming Lin, Halil Akin, Roshan Rao, Brian Hie, Zhongkai Zhu, Wenting Lu, Nikita Smetanin, Robert Verkuil, Ori Kabeli, Yaniv Shmueli, Allan dos Santos Costa, Maryam Fazel-Zarandi, Tom Sercu, Salvatore Candido, and Alexander Rives. Evolutionary-scale prediction of atomic-level protein structure with a language model. Science, 379(6637):1123–1130, 2023. doi: 10.1126/science.ade2574. URL https://www.science.org/doi/abs/10.1126/science.ade2574.

13. Kevin K. Yang, Nicolo Fusi, and Alex X. Lu. Convolutions are competitive with transformers for protein sequence pretraining. Cell Systems, 15:286–294.e2, 2024. doi: 10.1016/j.cels.2024.01.008.

14. Erik Nijkamp, Jeffrey Ruffolo, Eli N. Weinstein, Nikhil Naik, and Ali Madani. ProGen2: Exploring the boundaries of protein language models. arXiv, 2022.

15. Bo Chen, Xingyi Cheng, Pan Li, Yangli-ao Geng, Jing Gong, Shen Li, Zhilei Bei, Xu Tan, Boyan Wang, Xin Zeng, Chiming Liu, Aohan Zeng, Yuxiao Dong, Jie Tang, and Le Song. xTrimoPGLM: unified 100-billion-parameter pretrained transformer for deciphering the language of proteins. Nature Methods, 22(5): 1028–1039, May 2025. ISSN 1548-7105. doi: 10.1038/s41592-025-02636-z. URL https://doi.org/10.1038/s41592-025-02636-z.

16. Chloe Hsu, Robert Verkuil, Jason Liu, Zeming Lin, Brian Hie, Tom Sercu, Adam Lerer, and Alexander Rives. Learning inverse folding from millions of predicted structures. In Kamalika Chaudhuri, Stefanie Jegelka, L. Song, Csaba Szepesvari, Gang Niu, and Sivan Sabato, editors, Proceedings of the 39th International Conference on Machine Learning, volume 162 of Proceedings of Machine Learning Research, pages 8946–8970. PMLR, 17–23 Jul 2022. URL https://proceedings.mlr.press/v162/hsu22a.html.

17. Jin Su, Chenchen Han, Yuyang Zhou, Junjie Shan, Xibin Zhou, and Fajie Yuan. SaProt: Protein language modeling with structure-aware vocabulary. bioRxiv, 2023.

18. Pascal Notin, Lood Van Niekerk, Aaron W Kollasch, Daniel Ritter, Yarin Gal, and Debora S. Marks. TranceptEVE: Combining family-specific and family-agnostic models of protein sequences for improved fitness prediction. bioRxiv, 2022.

19. Mingchen Li, Yang Tan, Xinzhu Ma, Bozitao Zhong, Huiqun Yu, Ziyi Zhou, Wanli Ouyang, Bingxin Zhou, Pan Tan, and Liang Hong. ProSST: Protein language modeling with quantized structure and disentangled attention. In The Thirty-eighth Annual Conference on Neural Information Processing Systems, 2024.

20. Anthropic. Claude Opus 4.8 system card: the ProteinGym-Hard (PG-Hard) variant-ranking evaluation, 2026. URL https://www.anthropic.com/system-cards.

21. Alan F. Rubin, Hannah Gelman, Nathan Lucas, Sandra M. Bajjalieh, Anthony T. Papenfuss, Terence P. Speed, and Douglas M. Fowler. A statistical framework for analyzing deep mutational scanning data. Genome Biology, 18:150, 2017. doi: 10.1186/s13059-017-1272-5.

22. Benjamin J. Livesey and Joseph A. Marsh. Using deep mutational scanning to benchmark variant effect predictors and identify disease mutations. Molecular Systems Biology, 16(7):e9380, 2020. doi: 10.15252/msb.20199380.

23. Pascal Notin, Mafalda Dias, Jonathan Frazer, Javier Marchena-Hurtado, Aidan N Gomez, Debora Marks, and Yarin Gal. Tranception: Protein fitness prediction with autoregressive transformers and inference-time retrieval. In Kamalika Chaudhuri, Stefanie Jegelka, Le Song, Csaba Szepesvari, Gang Niu, and Sivan Sabato, editors, Proceedings of the 39th International Conference on Machine Learning, volume 162 of Proceedings of Machine Learning Research, pages 16990–17017. PMLR, 17–23 Jul 2022. URL https://proceedings.mlr.press/v162/notin22a.html.

24. Jiawei Gu, Xuhui Jiang, Zhichao Shi, Hexiang Tan, Xuehao Zhai, Chengjin Xu, Wei Li, Yinghan Shen, Shengjie Ma, Honghao Liu, Saizhuo Wang, Kun Zhang, Yuanzhuo Wang, Wen Gao, Lionel Ni, and Jian Guo. A survey on LLM-as-a-judge, 2025. URL https://arxiv.org/abs/2411.15594.

25. Google. What’s new in Gemini 3.5 Flash, 2026. URL https://ai.google.dev/gemini-api/docs/whats-new-gemini-3.5. Knowledge cutoff: January 2025; accessed 2 August 2026.

26. Anthropic. Claude models overview, 2026. URL https://platform.claude.com/docs/en/about-claude/models/overview. Claude Opus 4.8 reliable knowledge cutoff: January 2026; Claude Opus 5 reliable knowledge cutoff: May 2026; accessed 2 August 2026.

27. OpenAI. GPT-5.6 Sol model documentation, 2026. URL https://developers.openai.com/api/docs/models/gpt-5.6-sol. Knowledge cutoff: 16 February 2026; accessed 2 August 2026.

28. Nelson F. Liu, Kevin Lin, John Hewitt, Ashwin Paranjape, Michele Bevilacqua, Fabio Petroni, and Percy Liang. Lost in the middle: How language models use long contexts, 2023. arXiv:2307.03172.

29. Miles Turpin, Julian Michael, Ethan Perez, and Samuel R. Bowman. Language models don’t always say what they think: Unfaithful explanations in chain-of-thought prompting, 2023. URL https://arxiv.org/abs/2305.04388.

30. Tamera Lanham, Anna Chen, Ansh Radhakrishnan, Benoit Steiner, Carson Denison, Danny Hernandez, Dustin Li, Esin Durmus, Evan Hubinger, Jackson Kernion, Kamilė Lukošiūtė, Karina Nguyen, Newton Cheng, Nicholas Joseph, Nicholas Schiefer, Oliver Rausch, Robin Larson, Sam McCandlish, Sandipan Kundu, Saurav Kadavath, Shannon Yang, Thomas Henighan, Timothy Maxwell, Timothy Telleen-Lawton, Tristan Hume, Zac Hatfield-Dodds, Jared Kaplan, Jan Brauner, Samuel R. Bowman, and Ethan Perez. Measuring faithfulness in chain-of-thought reasoning, 2023. URL https://arxiv.org/abs/2307.13702.

31. Erna Sól Sigmarsdóttir, Vasileios Voutsinos, Kristoffer Enøe Johansson , Isa Kristin Henrichs, Alissa Buhrmann, Kresten Lindorff-Larsen, and Rasmus Hartmann-Petersen. Comprehensive variant effect map of parkin-mediated mitophagy in parkinson’s disease. bioRxiv, February 2026. doi: 10.64898/2026.02.09.704749. URL http://doi.org/10.64898/2026.02.09.704749.

32. Vahid Aslanzadeh, Sophie Glendinning, Yustika Sari, Marguerite Clarke, Christine J. Rodger, Xiaoyi Lin, Dipti Ranjan Lenka, Lucy Richardson, Jin Rui Liang, Teresa Kleinz, Duncan Sproul, Katja Lohmann, Christine Klein, Esther M. Sammler, Dario R. Alessi, Atul Kumar, Miratul M.K. Muqit, and Grzegorz Kudla. A functional genetic atlas of parkin resolves variants of uncertain significance and predicts parkinson’s disease age at onset. bioRxiv, February 2026. doi: 10.64898/2026.02.09.704817. URL http://doi.org/10.64898/2026.02.09.704817.

33. Mariano Martín and Benedetta Bolognesi. Systematic mutational mapping reveals optimal amyloid formation for ripk function. bioRxiv, February 2026. doi: 10.64898/2026.02.19.706818. URL https://doi.org/10.64898/2026.02.19.706818.

34. Samuel P. Berry, Camille B. Freedman, Debora S. Marks, and Rachelle Gaudet. Determinants of metal import and specificity in a bacterial transporter. bioRxiv, March 2026. doi: 10.64898/2026.03.30.714904. URL http://doi.org/10.64898/2026.03.30.714904.

35. Daeun Noh and Robert W. Newberry. Mutational scanning of α-synuclein using a clickable protein tag reveals determinants of membrane-induced aggregation. bioRxiv, April 2026. doi: 10.64898/2026.04.20.719647. URL http://doi.org/10.64898/2026.04.20.719647.

36. Connor Weyrick and Thomas Magliery. Next generation sequence counts for different E. coli cra mutant libraries after cell survival selection, 2026. URL https://datadryad.org/dataset/doi:10.5061/dryad.41ns1rnw3.

37. Changju Chun, Byeong-Kwon Sohn, Heesoo Ki, Ji Hye Jo, Hyoung-Tae An, Jinseo Park, Jiyu Lee, Sooyeon Choi, Jayeon Choi, Hyeyoung Cho, Seong Beom Lee, Booyoung Yu, Chae Young Lee, Ji Eun Kim, Yu-jin Ban, Young-Yoon Choi, Byoungsan Choi, Hongwon Lee, Junho Chung, Minkyung Baek, and Tae-Young Yoon. Landscape-scale navigation unlocks antibody cdr structural logic for ai-guided rescue and therapeutic optimization. bioRxiv, April 2026. doi: 10.64898/2026.04.21.719857. URL http://doi.org/10.64898/2026.04.21.719857.

38. Sabrina I. Giacometti, Nicolas Coudray, Rachel L. Redler, Gira Bhabha, and Damian C. Ekiert. Interactions of outer membrane lipoproteins P. aeruginosa pa3214 and E. coli pqic with their mce protein binding partners, pa3213 and pqib. bioRxiv, May 2026. doi: 10.64898/2026.05.09.724024. URL http://doi.org/10.64898/2026.05.09.724024.

39. Dana Cortade, James McLellan, Catherine Baranowski, Amanda Reider Apel, Peter J. Kelly, Aviv Spinner, Zach Sisson, Andi Dhroso, Corey Hudson, Simon dOelsnitz, Svetlana P. Ikonomova, and David Ross. Groq-seq function measurements for laci, ramr, and vanr transcription factor libraries — lmsf, March 2026. URL 10.5281/zenodo.19713339.

40. Siddharth Nimkar, Thu Nguyen, Deepti Karandur, Subu Subramanian, Michael E O’Donnell, and John Kuriyan. Redesign of energetically frustrated regions rescues function in defective t4 clamp loaders. bioRxiv, May 2026. doi: 10.64898/2026.05.08.723874. URL http://doi.org/10.64898/2026.05.08.723874.

41. Dana Cortade, James McLellan, Catherine Baranowski, Amanda Reider Apel, Peter J. Kelly, Aviv Spinner, Zach Sisson, Andi Dhroso, Corey Hudson, Anjali Chadha, Erika DeBenedictis, Svetlana P. Ikonomova, and David Ross. Groq-seq function measurements for tev protease ssvl and eppcr libraries., March 2026. URL 10.5281/zenodo.19713341.

42. Kai Fenzl, Linda H. Müller, Julia Kornienko, Maria Majchrowska, Nils Eikmeier, Benjamin Wehnert, Edukondalu Mullapudi, Brendan J. Floyd, Brian Clarke, Kristian Schweimer, Janosch Hennig, Daniel Schraivogel, Victoria N. Parikh, Matthias Wilmanns, and Lars M. Steinmetz. Multimodal phenotyping defines variant-to-function maps for rbm20 in dilated cardiomyopathy. bioRxiv, June 2026. doi: 10.64898/2026.06.10.730167. URL http://doi.org/10.64898/2026.06.10.730167.

43. Po-Yin Chen, Yung-Chih Chen, Chun-Hsiung Wang, Jia-Ying Su, Chien-Ling Lin, and See-Yeun Ting. Structural and functional principles of hcp-mediated antibacterial toxin delivery by the type vi secretion system. bioRxiv, June 2026. doi: 10.64898/2026.06.15.732508. URL http://doi.org/10.64898/2026.06.15.732508.

44. Bing Zhang, Pengbiao Xu, Yu Meng, Laure Gallay, François Lestelle, Hélène Morel, Marie-Louise Frémond, Bruno E. Correia, and Andrea Ablasser. The mutational landscape of sting-induced immunity. Nature, 655(8125):1271–1281, June 2026. ISSN 1476-4687. doi: 10.1038/s41586-026-10685-3. URL http://doi.org/10.1038/s41586-026-10685-3.

45. Warren van Loggerenberg, Haotian Zhang, Vignesh Senguttuvan, Michael J. Chambers, Mailoan Panchalingam, Alireza Rasoulzadeh, Anna Axakova, Marinella Gebbia, Robert J. Desnick, Bruce Wang, Caroline Schmitt, Laurent Gouya, Jordi To-Figueras, Alexander Wahl, Ivet Bahar, Pemra Doruker, and Frederick P. Roth. Environment-dependent landscapes of coding variant impacts on coproporphyrinogen oxidase. bioRxiv, July 2026. doi: 10.64898/2026.07.02.735896. URL http://doi.org/10.64898/2026.07.02.735896.

46. Jerry C. Dinan, James W. McCormick, Rishi Soni, Samuel Thompson, and Kimberly A. Reynolds. Protein surface site determines the evolutionary accessibility of allosteric regulation. bioRxiv, July 2026. doi: 10.64898/2026.07.02.735819. URL http://doi.org/10.64898/2026.07.02.735819.

47. Priyanka Madhu, Caroline Benz, Leandro Simonetti, Matthew J. Winters, Mythili S. Subbanna, Johanna Kliche, Lidia Gomez-Lucas, Hazem M Kotb, Izabella Krystkowiak, Dmitri Segal, William T. P. Darling, Sven Larsen-Ledet, Maximilian Vieler, Ana Zupancic, Aimiliani Konstantinou, Filip Mihalic, Julia K Varga, Livia Pagano, Susanne Lüchow, Andreas Kraemer, Lachlan Ellingboe, Kristof Görlitz, Germanna L. Righetto, Christin Kossmann, Ruisheng Xiong, Ora Schueler-Furman, Levon Halabelian, Vijayaratnam Santhakumar, Cheryl Arrowsmith, Stefan Knapp, Mate Erdelyi, Amelie Stein, Renaud Vincentelli, Peter M. Pryciak, Norman E Davey, and Ylva Ivarsson. An atlas of short linear motif-mediated human protein-protein interactions. bioRxiv, July 2026. doi: 10.64898/2026.07.03.735260. URL http://doi.org/10.64898/2026.07.03.735260.

48. John T. Lazar, Evan Komp, Irene Martinez, Kyle S. Zolkin, Pascal Notin, Samer Saleh, Grant M. Landwehr, Kyung G. Kim, Anru Tian, Benjamin A. Shapero, Ashty S. Karim, Debora S. Marks, Gregg T. Beckham, and Michael C. Jewett. Machine learning guided cell-free expression maps the biochemical landscape of carbonic anhydrase. bioRxiv, July 2026. doi: 10.64898/2026.07.07.736810. URL http://doi.org/10.64898/2026.07.07.736810.

49. NCBI Gene Expression Omnibus. Genotype-phenotype single-cell transcriptomics for massive parallel assessment of genetic variants, 2026. URL https://www.ncbi.nlm.nih.gov/geo/query/acc.cgi?acc=GSE311874. GEO accession GSE311874; public 10 July 2026.

